# Very high density EEG elucidates spatiotemporal aspects of early visual processing

**DOI:** 10.1101/172353

**Authors:** Amanda K Robinson, Praveen Venkatesh, Matthew J. Boring, Michael J. Tarr, Pulkit Grover, Marlene Behrmann

**Author notes:** Shared contribution (Pulkit Grover, Marlene Behrmann). Corresponding author, Amanda K. Robinson, Department of Psychology, Carnegie Mellon University, Pittsburgh PA 15213.

## Abstract

Standard human EEG systems based on spatial Nyquist estimates suggest that 20-30 mm electrode spacing suffices to capture neural signals on the scalp, but recent studies posit that increasing sensor density can provide higher resolution neural information. Here, we compared “super-Nyquist” density EEG (“SND”) with Nyquist density (“ND”) arrays for assessing the spatiotemporal aspects of early visual processing. EEG was measured from 128 electrodes arranged over occipitotemporal brain regions (14 mm spacing) while participants viewed flickering checkerboard stimuli. Analyses compared SND with ND-equivalent subsets of the same electrodes. Frequency-tagged stimuli were classified more accurately with SND than ND arrays in both the time and the frequency domains. Representational similarity analysis revealed that a computational model of V1 correlated more highly with the SND than the ND array. Overall, SND EEG captured more neural information from visual cortex, arguing for increased development of this approach in basic and translational neuroscience.

**Abbreviations:**

SND: “super-Nyquist” density EEG (smaller than 20-30 mm electrode spacing)
ND: Nyquist density EEG (20-30 mm electrode spacing)

## Introduction

As a non-invasive neuroimaging method that is almost unrivalled in its temporal precision, electroencephalography (EEG) is extremely useful in studying the time course of neural processes. EEG also has the advantage of being less expensive and easier to implement than the similarly time-precise method of magnetoencephalography (MEG). However, relative to functional magnetic resonance imaging (fMRI) and, to a lesser extent, MEG, EEG signals are believed to yield relatively low spatial resolution. This is thought to be due to the disproportionate decay of high-spatial frequencies (which carry high resolution information) during volume conduction from electrical sources in the brain to electrodes on the scalp. At the same time, the concrete limits of EEG’s spatial resolution are not well understood, and a critical question remains unanswered: does increasing the density of EEG electrodes enable the extraction of high-resolution spatial information? This study seeks to provide an answer to this question through an experiment designed to test whether, in human early visual cortex, high-density EEG can capture higher-resolution spatial neural information as compared to present-day, standard low-density systems.

Prior results estimated the “spatial Nyquist rate” of EEG and concluded that inter-electrode distances of about 20-30 mm suffice to extract the maximum possible resolution from EEG signals (Srinivasan, Nunez, & Silberstein, 1998; Srinivasan, Tucker, & Murias, 1998). While these results have led to continued skepticism regarding the utility of higher-density EEG, recent theoretical work establishes that these approximations of spatial Nyquist rates severely underestimate the required number of sensors (Grover & Venkatesh, 2017). Conceptually, the spatial Nyquist rate of EEG specifies the number of electrodes needed so that the EEG signal can be recovered at any point on the scalp by appropriately interpolating the electrodes’ values. A frequently-used approximation of the Nyquist rate guarantees that the sampled signal can recover a large percentage (e.g., 95%) of the EEG signal’s energy (only 5% error in mean-square sense of the signal on the scalp) (Srinivasan, Nunez, et al., 1998; Srinivasan, Tucker, et al., 1998). This density, which corresponds to 20-30 mm inter-electrode separation, is referred to as the Nyquist Density in this paper, and is the basis for the design of standard EEG systems. Nyquist-density grids do sample scalp signals to within a small error, but have the unfortunate disadvantage of being unable to capture high-resolution signals; the disproportionate decay of high spatial frequencies leads to low amplitude high-resolution signal at the scalp, which cannot be recovered with Nyquist-density grids (Grover & Venkatesh, 2017). However, very often, the goal of EEG sensing is not to recover the high amplitude, low spatial frequency signal on the scalp itself, but instead to detect subtle differences in the spatial patterns of activity on the scalp, or to make inferences about the signal source *inside the brain.* For such inferences, the high-resolution (but low-amplitude) information can be critical, and higher density sampling that recovers this information could benefit the inference immensely.

Indeed, there is experimental evidence that supports the claim that denser EEG systems can recover more information: Freeman et al. (2003) and Petrov et al. (2014) used high-density EEG configurations to estimate that 10 mm inter-electrode spacing provides additional neural information. However, both Freeman et al. (2003) and Petrov et al. (2014) included limitations that leave open the question as to whether these studies do demonstrate the utility of going beyond EEG Nyquist rate estimates (Srinivasan, Nunez, et al., 1998; Srinivasan, Tucker, et al., 1998). First, both studies used electrode arrays that have small spatial coverage: Freeman et al. used a *linear* array of 64 electrodes, and Petrov et al. used a small 4x4 grid of electrodes. This limited scalp coverage leaves open the possibility that lower density but higher coverage grids could still recover the signals sampled at high-density locations. Additionally, the experimental protocols employed were limited in their scope; Freeman et al. estimated spatial and temporal power spectral densities using a resting state design, but did not attempt to determine whether the additional neural information gathered with ultrahigh density electrodes was meaningful. Petrov et al. showed that word/non-word classification was more accurate with an ultra-high density grid over parietal cortex, but made no concrete neural predictions as to why higher density sampling was beneficial in this case. In contrast, the goal of the current research is to compare neural information obtained by a high-coverage, high-density (and hence *Super*-Nyquist density) EEG system with that from standard Nyquist density counterparts in an experimental paradigm utilizing carefully-controlled stimuli which have expected neural responses within early sensory areas of the brain.

The specific domain in which we conduct this comparison is that of human low-level vision. Humans are remarkably adept at recognizing and identifying a multitude of different visual stimuli. As part of this process, early visual cortex responds to low-level visual information such as edges, orientation, contrast and spatial frequency and does so in a retinotopic manner, such that responses approximate a polar coordinate system (Tootell et al., 1998). More importantly for our purposes, the temporal and spatial aspects of the neural processes involved in low-level visual perception have been well characterized (Francesco Di Russo et al., 2005, 2007; Engel et al., 1994). For this reason, early vision offers a unique opportunity to investigate how well EEG can harness meaningful spatial information from the scalp.

Our current study investigated the spatial and temporal mechanisms of visual processing using high- versus low-density EEG. To distinguish between nomenclature variations with respect to EEG density, we refer to the very high density EEG array as “super-Nyquist density” (SND), and the lower-density EEG array comparable to current systems as “Nyquist density” (ND). To accomplish these two variations, we modified an EEG head cap from a 128-electrode Biosemi system, increasing sensor density 2- to 3fold over occipitotemporal regions. To our knowledge, this is the highest density EEG system for this extent of coverage (Fig. 1a) to date. In our experiment, we designed visual stimuli that had low-, medium- and high-spatial-frequency content. Due to the small receptive field sizes of neurons in early visual cortex, we predicted that these stimuli would produce neural responses that reflected their relative spatial-frequency characteristics. By utilizing a visual paradigm designed to elicit neural responses with differing spatial frequencies in the brain, we examined how SND EEG improves spatial capture of visual neural information at the scalp. While a conventional spatial Nyquist rate analysis would predict no benefits of the increased sensor density, our results show that SND EEG provides substantial and significant benefits over standard EEG sensor densities. Such results are extremely promising for future applications of high-density EEG in both basic and translational settings.

**Figure 1.**
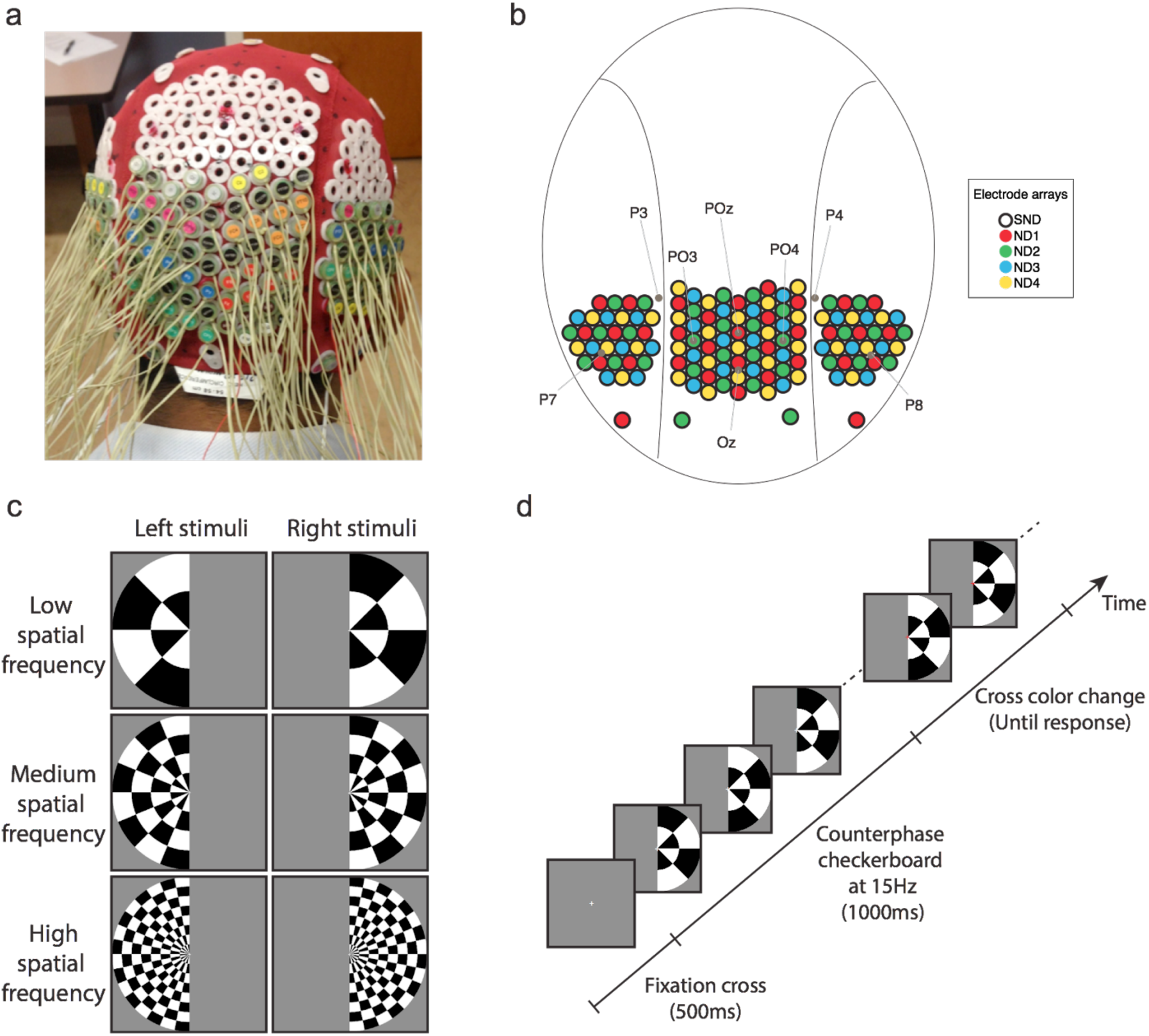
Experimental design. (a) Modified EEG head cap with high-density electrode configuration (b) Map of electrode configuration, showing the SND layout and subset of electrodes designated for each ND array. Some standard electrode locations are marked for reference. (c) Stimuli and design of the experiment: stimuli were low, medium or high spatial frequency checkerboards presented in the left or right visual field. d) An example trial. A central gray fixation cross was presented for 500 ms, then the checkerboard stimulus counter-phased at 15 Hz. After 1000 ms, the cross became red or green, at which time participants discriminated its color and responded via a key-press.

## Materials and Methods

### Participants

This study was approved by the Institutional Review Board of Carnegie Mellon University (CMU). Informed consent was obtained from all participants. Participants were 20 adults recruited from CMU (2 females). They were compensated US $50. All participants tested had head sizes approximately 590-620 mm circumference in order to fit the SND EEG cap. Four participants were excluded due to excess eye movements or poor data quality, leaving *N* =16.

### SND and ND EEG recording

“Super-Nyquist” (very high density) EEG was measured on participants using a standard 128-electrode BioSemi Active Two system (BioSemi, Amsterdam, Netherlands) with custom electrode configuration over occipitotemporal brain regions. The 128 electrodes, approximately 14 mm apart, were arranged on a nylon head cap in a high-density configuration over the back of the head (Figure 1a,b). Four ND configurations were chosen as subsets of the 128 SND electrodes (Figure 1b) measured at the same time. For the ND arrays, electrode distance was approximately 28 mm, which is equivalent to current commercial 64-128 electrode EEG systems. SND EEG thus contained identical information to the combination of the four ND arrays.

Continuous EEG data were recorded and digitized at a 512 Hz sample rate with 24-bit A/D conversion. During recording, all scalp electrodes were referenced to the standard BioSemi reference electrodes. Eye movements were monitored using bipolar horizontal EOG electrodes placed at the outer canthi of each eye and bipolar vertical EOG electrodes placed above and below the left eye.

### Design and Stimuli

Experimental stimuli consisted of checkerboards of low, medium or high spatial frequency with radius 28 degrees of arc (Figure 1c). The spatial frequencies were .036, .071 and .143 cycles per degree radially (equivalent to 1, 2 and 4 wavelengths of alternating black/white tiles), and .011, .022 and .044 cycles per angular degree (equivalent to 2, 4 and 8 wavelengths in a semicircle), respectively. The patterns of the checkerboards reversed at a frequency of 15 Hz for 1000 ms. The stimuli appeared in either the left or right visual field. Spatial frequency and hemifield were sampled with equal probability in each block of trials. Following the 500 ms initial fixation and 1000 ms checkerboard presentation, the central fixation cross became red or green (Figure 1d). Participants performed a simple orthogonal task, namely to discriminate the color of the fixation cross using key presses. No feedback was given regarding accuracy.

The Psychophysics Toolbox in Matlab was used to present the visual stimuli on a 24-inch LCD monitor with 60 Hz refresh rate. Across ten blocks of trials there were 100 repeats of each of the 6 stimulus types, resulting in 600 trials for the whole experiment. A small break between blocks was offered to observers.

### EEG preprocessing

EEG data analysis was performed offline using EEGlab (Delorme & Makeig, 2004). The data were preprocessed using the PREP pipeline (Bigdely-shamlo, Mullen, Kothe, Su, & Robbins, 2015): data were temporarily high pass filtered at 1 Hz, line noise was filtered and average reference was applied. No electrodes were interpolated. For analysis, data were subjected to low-pass (0.1 Hz) and high-pass (100 Hz) zero-phase filters. Epochs were constructed relative to first checkerboard onset and baseline corrected for 200 ms prior to onset. EEG data were then subsampled to 256 Hz using the decimate function in Matlab.

### Analyses

Three main analyses were performed to assess the utility of high-density EEG: (1) decoding stimulus representations across the time-course of a trial, (2) decoding stimulus representations in the time-frequency domain, and (3) representational similarity analyses (RSA) to assess how well the neural data fit a model of primary visual cortex (V1) activity. Analyses were performed using all 128 very-high-density electrodes (SND), as well as four subsets of the same electrodes, each equivalent to standard density electrode arrays (ND1-ND4; see Figure 1b).

### Time course decoding

A support vector machine (SVM) classifier was used to decode the neural activity evoked by the six different stimuli. For each participant, trials were excluded if they had any horizontal eye movements or contained more than 10 channels exceeding 150 uV over the course of the trial. From the remaining included trials, 80 trials per stimulus condition were chosen randomly for decoding analyses to equate trial numbers across participants. To improve decoding performance, two trials were averaged before classification: the mean was taken for two trials (approximately one from blocks 1-5 and one from blocks 6-10) to reduce non-stimulus related EEG noise (Grootswagers, Wardle, & Carlson, 2017). This resulted in 40 averaged pseudotrials per stimulus condition (240 trials in total) used for decoding. Trial averaging did not change the trend of the results (SND>ND), but improved decoding performance in general.

For all analyses, six-way classification was performed for all stimulus types (frequency X field). Essentially, multiclass SVM conducted all possible pairwise classification tests using 10-fold cross-validation. Data for each electrode were *z*-scored across all trials to standardize electrodes before classification. Principal components analysis (PCA) was performed to reduce data dimensionality. At each fold of cross-validation, PCA was performed on the training data and weights were applied to the test data (Grootswagers et al., 2017). PCA components accounting for 99% of the variance in the training data were used as features for classification. Analyses were conducted for each participant, time point, and for the SND and ND electrode arrays.

### Frequency domain decoding

A fourier transform was applied to the 240 averaged pseudotrials from 0-1000 ms (256 samples) using the Letswave 6 toolbox. The amplitude at 15 Hz was selected for each electrode, and *z*-scored across trials. These data were then fed to the classifier. The same parameters were used for decoding as in the time course data; namely, six-way SVM with 10-fold cross-validation was conducted on the *z*-scored data and feature selection involved PCA at each fold. Features were PCA components accounting for 10, 20, 30, 40, 50, 60, 70, 80, 90 and 99% of the variance across trials.

### Representational similarity analyses

#### HMAX RDM

A hierarchical computational model of object representation (HMAX; Riesenhuber & Poggio, 2000) was used to determine the dissimilarity between experimental stimuli in terms of their expected processing by visual cortex. The six visual stimuli were fed through the S1 layer of HMAX, which was designed to reflect retinotopically-organized simple cell response properties in V1 (Serre, Oliva, & Poggio, 2007). A representational dissimilarity matrix (RDM) was populated by calculating Euclidean distance between S1 outputs for each pair of stimuli.

#### EEG RDM

For each stimulus, a mean event-related potential (ERP) trace was constructed per electrode. Trials were excluded before averaging if there were eye movements or more than 25 electrodes exceeded 150 uV from 0-300 ms. Electrodes were removed if they would have normally been interpolated (as assessed by the PREP pipeline), if they exceeded the voltage threshold on more than half the trials, or if the standard deviation of their grand average ERP trace was higher than 5 across the trial. The mean numbers of electrodes used for analyses were 120.44 for SND, 28.13 for ND1, 30.82 for ND2, 30.88 for ND3 and 30.63 for ND4. At each time point, Euclidean distance was calculated for each pair of stimuli across electrodes to populate the RDM. To normalize Euclidean distance values across electrode sets, a correction was made to account for the number of electrodes included in the analysis.

#### RSA between HMAX and EEG

Spearman correlation was calculated between the HMAX RDM and EEG RDM at each time point per participant and per electrode array (SND, ND1-4). Correlation values were Fisher-transformed before averaging across participants.

### Statistical analysis

For each analysis, performance using the SND electrodes was compared with the four ND electrode arrays using two-tailed pairwise *t*-tests. Uncorrected *p* values are presented. Effect sizes were calculated for all statistical comparisons; generalised eta-squared values (*η*_g_^2^) were calculated for ANOVA results, and Hedges’ *g*_av_ was calculated for paired samples *t*-tests (Bakeman, 2005; Lakens, 2013).

## Results

Based on the theoretical analyses that challenge the approximate Nyquist analysis (Grover & Venkatesh, 2017), we predicted that SND electrodes would capture more neural information than ND, and therefore expected to see SND outperform ND in all analyses. Furthermore, high spatial frequency stimuli were expected to produce neural patterns with the highest spatial frequency, so we expected to see a greater SND advantage for the high spatial frequency stimuli than low spatial frequency stimuli.

### Time course decoding

In the time domain, SND EEG classified stimulus representations significantly better than ND EEG. At each time point across the course of a trial, a linear support vector machine (SVM) classifier was used to decode the neural activity evoked by the different stimuli. Six-way classification was performed for all stimuli (low, medium and high spatial frequency stimuli in left and right visual fields). As shown in Figure 3, decoding was at chance (16.66%) during the baseline period. Significant decoding was observed from approximately 80 ms after image onset, a time period associated with early visual processing. A peak in decoding accuracy was observed at approximately 105 ms, which corresponds with the P1 or P100 event-related potential (ERP) component in typical EEG, and has been associated with early visual cortex activity (Di Russo, Martinez, & Hillyard, 2003). These results suggest that significant decoding relied upon early visual cortex representations of low-level visual features that differed across the six stimuli.

At the time of the peak in decoding accuracy, determined as the 10 ms time windows encompassing each individual’s peak decoding performance within the 80-140 ms time window (Figure 2c and 2d show example participant traces), a one-way ANOVA revealed a significant effect of electrode array, *F*_4,60_ = 10.06, *p* < .001, *η_G_*^2^ = .046, with SND outperforming all ND arrays, *ts* > 3.27, *ps* < .006, *g*_av_ > .333. ND1 also resulted in significantly lower decoding accuracy than ND2, ND3 and ND4, *ts* > 2.47, *ps* < .026, *g*_av_ > .161, but there were no differences across ND2, ND3 and ND4, *ts* < 1.37, *ps* > .190, *g*_av_ < .121.

**Figure 2.**
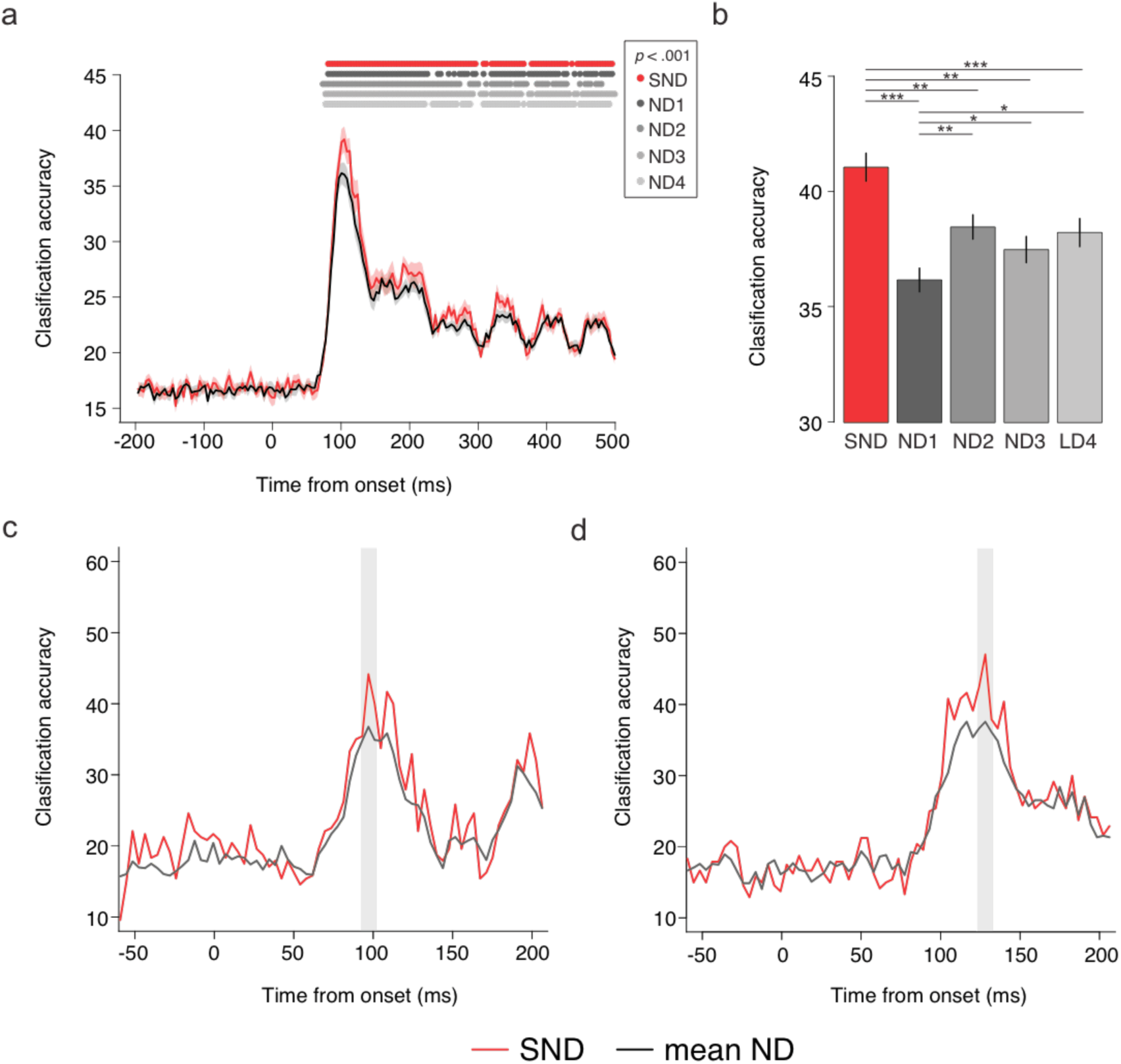
Decoding results. (a) Mean accuracy for 6-way classification, performed for each time point for the SND and mean of ND electrode arrays. Shaded area depicts standard error of the mean corrected for within subjects designs (Cousineau & O’Brien, 2014). Dots depict time points with significant group-wise decoding per array, *p* < .001. (b) Mean decoding accuracy per electrode array, 10 ms around each individual’s first peak in decoding. SND outperformed all ND arrays. (c) Example decoding trace for one participant, with peak at 92-102 ms. (d) Example decoding trace for one participant, with peak at 123-133 ms.

These results indicate that SND electrodes capture more neural information than ND EEG arrays. As expected, the decoding advantage of SND EEG was evident at the peak in classification; this is likely to be the peak time of the neural stimulus representations in early visual cortex, and thus include neural activity with high spatial frequency. Analysis of the classification confusion matrices revealed that the SND decoding advantage was greater for high than low and medium spatial frequency stimuli (see Supplemental Material), supporting the hypothesis that SND EEG captures more high spatial frequency neural information at the scalp. Variation across the ND electrode arrays, all of which had the same electrode density and approximate scalp coverage, further suggests that certain ND configurations provide differential neural information, and that information relevant to classification is present on the scalp at distances closer than the spacing of ND electrodes.

### Frequency domain decoding

One inherent drawback of the time domain decoding analyses is that significant decoding could be due to differences in spatial *or* temporal aspects of stimulus processing. In other words, the superiority of SND in time course decoding might be caused by heightened sensitivity to subtle changes in the latency of responses for the different stimuli, rather than from the spatial pattern of neural stimulus representations. To rule out this possibility and assess the advantage of SND EEG in the frequency domain, we capitalized on the frequency tagging protocol of the experiment by decoding stimulus responses in time-frequency space, using what has been termed the steady state visual evoked potential (SSVEP) (Regan, 1966). A fast Fourier transform was applied to each averaged pseudotrial 0-1000 ms from stimulus onset, and the amplitude at 15 Hz (the stimulus flicker frequency) for each electrode was fed to the classifier. This analysis established both whether the spatial SSVEP information captured by EEG can reliably distinguish between stimuli, and whether SND EEG results in greater accuracy than ND. Additionally, we explored how classification varied when different numbers of features were fed to the classifier. To reduce the overall dimensionality of our data, features were PCA components accounting for 10, 20, 30, 40, 50, 60, 70, 80, 90 and 99 percent of the variance in the dataset (Figure 3).

**Figure 3.**
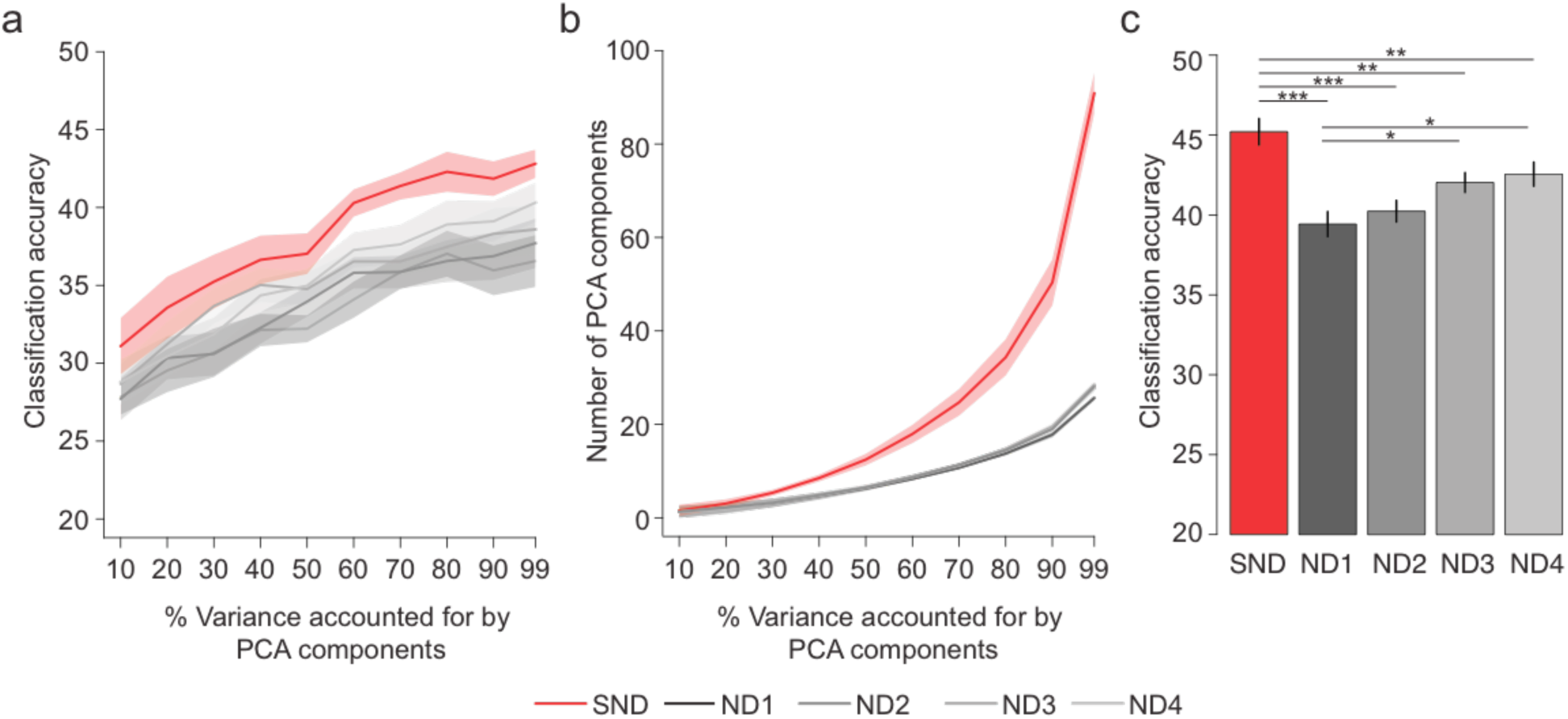
Decoding accuracy for frequency domain data. (a) Classification accuracy as function of the number of PCA components fed to the classifier per electrode array. SND had the best decoding accuracy for every level of PCA components. (b) Number of PCA components as a function of the percent variance accounted for by the PCA components. These were the number of features used for classification. (c) Decoding accuracy for frequency domain data, using maximum decoding accuracy per electrode array per participant. SND had significantly larger decoding accuracy than all ND arrays. ^∗^*p* < .05, ^∗∗^*p* < .01, ^∗∗∗^*p* < .001.

SND outperformed all ND configurations regardless of the number of features (PCA components) fed to the classifier (Figure 3a). A 5 x 10 ANOVA with factors of electrode array and number of features revealed a significant main effect of electrode array, *F*_4,60_ = 9.52, *p* < .001, *η*_G_^2^ = .154, with SND outperforming all ND arrays, *ts* > 3.38, *ps* < .002, *g*_av_ > .292, but no significant difference across the four ND arrays, *ts* < 2.056, *ps* > .057, g_av_ < .233. There was also a significant main effect of features, *F*_1,15_ = 15.14, *p* = .001, *η*_G_^2^ = .391, such that more features resulted in better classification. The interaction between electrode array and features did not reach significance, *F*_4,60_ = 1.70, *p* = .161, *η*_G_^2^ = .009. Importantly, even when the highest classifier accuracy was chosen per set for each participant, SND still outperformed ND by 3-6%, *ts* > 3.05, *ps* < .009, *g*_av_ > .237 (Figure 4c). This analysis also revealed a small, but significant, difference between ND1 and ND3, and between ND1 and ND4, *ts* > 2.51, *ps* < .025, *g*_av_ > .233. There were no significant differences across the other ND arrays, *ts* < 2.08, *ps* > .055, *g*_av_ < .216.

**Figure 4.**
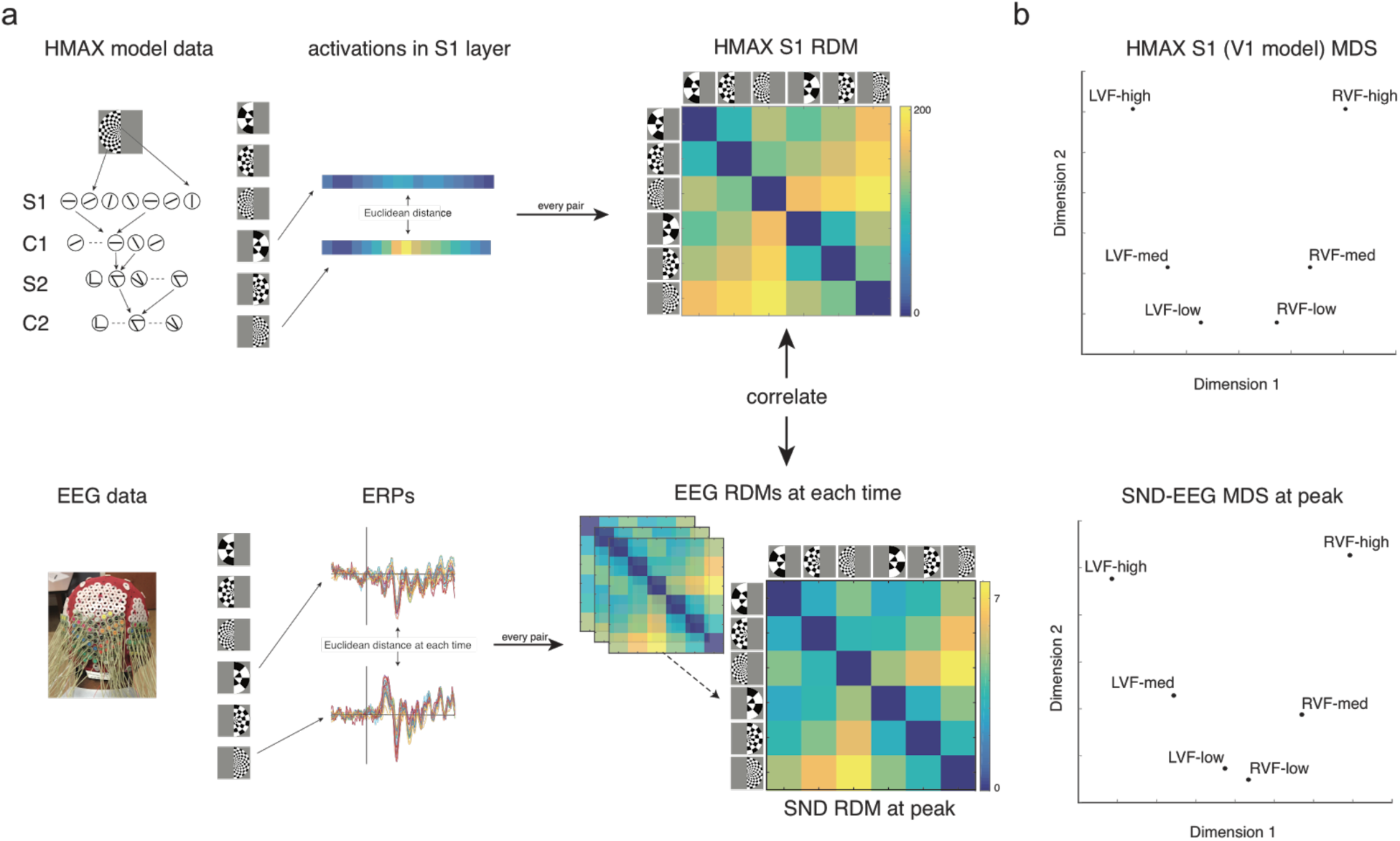
Representation similarity analysis. (a) RSA pipeline. Dissimilarity matrices (RDMs) were computed based on Euclidean distance for each pair of stimuli using HMAX S1 (V1 model) and EEG data. Correlations were then performed between the HMAX model RDM and the EEG RDM at each time point. (b) Multidimensional scaling (MDS) for HMAX model and SND EEG RDM at the first peak. The separation of stimuli was very similar for SND EEG at the first peak and HMAX S1. LVF = left visual field stimulus, RVF = right visual field stimulus. Low, med and high refer to spatial frequency of stimuli.

Our frequency domain analyses complement our time course analyses in that SND EEG results in better classification of neural data than ND electrode arrays with the same scalp coverage. These analyses also reveal some variability in decoding performance across ND configurations, suggesting that electrode positioning might influence the degree of neural information capture at the scalp. SND superiority for SSVEP decoding reinforces the observation made from time-domain decoding: SND EEG harnesses higher spatial frequency neural information, improving spatial resolution of neural signals.

### Representational Similarity Analyses

EEG is a particularly good technique for assessing the time course of neural processing, but current approaches lack detail in spatial resolution. The time course and SSVEP decoding results indicate that EEG can discriminate different visual responses, but it remains unclear how the stimulus representations captured by EEG relate to early visual cortex activity. Addressing this question, the simple stimuli and SND electrode configuration used in our study also offer an opportunity to assess how well EEG captures spatial aspects of neural representations, for example the retinotopic nature of V1 responses. We used representational similarity analysis (RSA), a technique that computes a measure of dissimilarity for different neural representations of the stimuli and compares the resultant pattern against a model of neural activity at different stages of processing (Kriegeskorte, Mur, & Bandettini, 2008).

In our RSA analysis, we were interested in the extent to which the neural dissimilarity between stimuli (as measured by EEG) reflects differences in the low-level visual features of the stimuli, which are likely to differ in their spatial patterns of neural representation. To accomplish this, we compared two measures of stimulus dissimilarity: (1) one based on HMAX, a model designed to represent activity in visual cortex (Riesenhuber & Poggio, 2000), and (2) one based on evoked EEG activity in response to the different stimuli at each time point. To compare these two representational dissimilarity measures, Spearman correlations were computed between the HMAX and EEG dissimilarity results for the 15 unique pairwise stimulus values. A positive correlation thus indicated that stimulus representations measured by EEG were related in the same way as well-characterized responses in early visual cortex, suggesting that EEG was directly measuring early visual cortical activity. RSA correlations were compared between SND and ND EEG to assess the hypothesis that SND EEG would capture more spatially relevant visual activity.

To measure the expected stimulus pattern dissimilarity based on known responses in visual cortex, we ran the stimulus images through HMAX and obtained values from level S1, a model designed to reflect simple cell activity in primate primary visual cortex (V1), discriminating low level features in a retinotopic manner (Riesenhuber & Poggio, 2000). Euclidean distance was then calculated for all pairwise comparisons of the six stimuli to construct the model representational dissimilarity matrix (RDM; Figure 4a). The HMAX S1 RDM revealed that the difference between the medium and high spatial frequency stimuli was higher than the difference between low and medium spatial frequencies (Figure 4b). Furthermore, the difference between left and right visual field stimuli was larger for high spatial frequency than low spatial frequency stimuli. Multidimensional scaling (MDS; Figure 4b) showed a clear delineation between stimuli based on their visual field and spatial frequency.

To construct the EEG dissimilarity measure, we calculated Euclidean distances for the ERP voltage across electrodes for all pairwise comparisons of stimuli at each time point from image onset (see Supplementary Material). In accordance with the decoding results, it was clear that the greatest separation between conditions (that is, a peak in Euclidean distance) occurred approximately 100 ms after stimulus onset. Euclidean distance was normalized for the number of electrodes included in the analysis, so there was no significant difference between the SND and ND arrays in terms of absolute dissimilarity values at the 100 ms peak. The EEG RDMs at the peak appeared to show the same pattern as the RDM from the HMAX values, and MDS on these results revealed a similar separation between visual field and spatial frequency of the stimuli (Figure 4a and 4b).

Figure 5 shows the most informative RSA results: the correlation between HMAX and EEG RDMs across time. As expected, the correlation values were approximately zero from the baseline period up to 50 ms following image onset, and did not differ between SND and ND at this time. Correlation values peaked around 100 ms, reflecting early visual cortical activity (Spearman correlation at the peak was *r* = .79). This time point also exhibited a dissociation between the electrode arrays.

**Figure 5.**
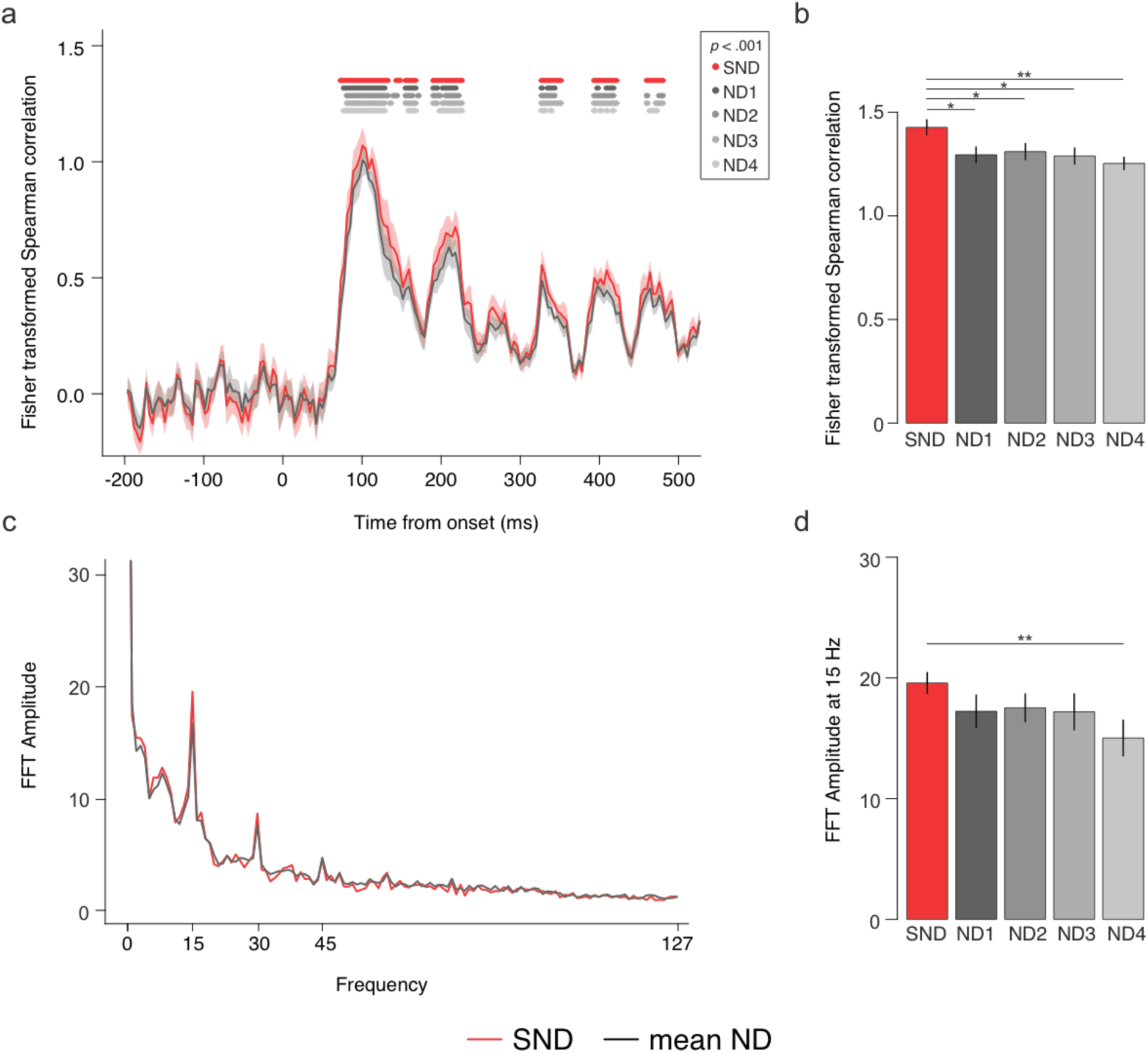
(a) Correlation between HMAX and EEG stimulus dissimilarity across time for each electrode array (high density and mean low density). Dots signify significant groupwise correlations per array. (b) Mean correlation for each electrode array, 10 ms around each individual’s first peak within the time period 80-140 ms. SND was significantly more correlated than ND. (c) Fourier transform of RSA correlation values 0-1000 ms. (d) Mean 15 Hz signal in correlation values per electrode array. ^∗^*p* < .05, ^∗∗^*p* < .01.

At the time of maximum correlation, there was a significant effect of electrode array, *F*_4,60_ = 3.26, *p* = .018, *η*_g_^2^ = .033. SND was significantly higher than all ND arrays, *ts* > 2.30, *ps* < .036, *g*_av_ > .374 (Figure 5b). There were no significant differences across ND arrays, *ts* < 1.33, *ps* > .203, *g*_av_ < .179. Noise ceiling calculations for each electrode array revealed that both SND and ND RSA correlations reached the lower bound of the noise ceiling at this peak, indicating that the HMAX S1 model is a very good explanation for EEG data during this time period (see Supplementary Material). Importantly, however, the SND array had a higher upper bound of the noise ceiling, indicating the SND array offered greater ability to calculate the true representational similarity model at the group level. Other layers of HMAX were also tested against the EEG RDMs. For the first peak, the S1 layer of HMAX had the highest correlation. C1 and S2, subsequent layers that also represent low-level aspects of stimuli, both had lower, but significant correlations at this peak. The C2 layer did not reach significance at any time point at the group level, although a small peak was evident from approximately 120 ms, which is in line with the hierarchical nature of the HMAX model. C2 is meant to represent later, view-invariant responses, so would be expected to correlate at a later time point with the EEG data (see Supplementary Material).

Overall, these RSA analyses provide strong evidence that the stimulus representations measured by EEG contained meaningful spatial neural activity evoked from V1. Specifically, the EEG RDM was significantly correlated with the HMAX V1 model RDM at approximately 100 ms, providing strong evidence that EEG captured retinotopic neural information in V1. Furthermore, the SND array RDM had a stronger correlation than the ND arrays, indicating that it captured more stimulus-relevant spatial activity. This is striking because the absolute values of the EEG RDM did not vary between SND and ND electrode arrays, that is, the EEG data from both arrays produced the same mean separation across stimulus pairs. Hence, the increased correlation of SND between EEG- and HMAX-RDM values must be due to SND having a more accurate representation of the separation between stimulus pairs. Essentially, while both SND and ND electrode arrays appear to characterize early visual cortical activity, the SND advantage suggests that additional scalp electrodes enable better resolution of relevant neural signals.

Interestingly, while groupwise correlation values stayed higher than the baseline for most of the trial period, there were periodic fluctuations at 15 Hz, the same frequency as the image presentation. To determine how well the SSVEP signal was represented by the EEG-HMAX RSA, we conducted a Fourier transform on the correlation values for the trial period 0-1000 ms. As can be see in Figure 5c, the Fourier plot revealed peaks at 15 Hz and its harmonics, indicative that the periodic EEG activity was characteristic of periodic early visual cortex activity at the visual stimulation frequency. Furthermore, this 15 Hz signal was significantly higher for the SND than the ND4 array, *t* = 3.00 *p* = .009, *g*_av_ = .344. SND was higher than ND1, ND2 and ND3, but this did not reach significance, *ts* > 1.415, *ps* < .178, *g*_av_ > .141. There were no significant differences across ND arrays, *ts* < 1.17, *ps* > .260, *g*_av_ < .179. Although S1 had the highest 15 Hz signal and was thus the best explanation of the EEG data, analysis of the periodic signal from C1, S2 and C2 layers of HMAX revealed a 15 Hz signal from all layers, suggesting that visual processing progressed from simple cell activity in this paradigm, and that EEG could detect these separate but related processes. This result demonstrates that EEG can capture periodic activity associated with processing in early visual cortex, and indicates that SND EEG captures more relevant periodic stimulus information than ND-EEG, for at least one of the ND arrays.

Overall, these results suggest that our RSA correlation measure, based on the S1 layer of the HMAX model, captures meaningful spatial patterns of neural activity measured at the scalp, specifically relating EEG activity to well-characterized retinotopic simple cell responses in V1. In addition, it is clear that SND EEG can capture more spatial information than equivalent ND-EEG, and this additional neural information offers greater insight into the spatiotemporal aspects of early visual processing.

## Discussion

In this study, in the first time, to our knowledge, systematically comparing an SND EEG system of such a high density and extent of scalp coverage to equivalent ND arrays, we found that “super-Nyquist” density EEG captures considerably more spatial neural information at the scalp than current Nyquist density systems. Across three separate analyses, SND EEG represented neural activity in human early visual cortex in a more robust and meaningful manner than ND electrode arrays with the same scalp coverage. SND EEG outperformed ND-EEG in decoding between six different visual stimuli varying in two low-level visual features (spatial frequency and visual field) in both the time and the frequency domains. Furthermore, SND EEG patterns of neural activity were found to better approximate V1 representations, defined independently using a well-established computational model of cortical visual processing. This SND advantage indicates that higher density electrodes can capture more fine-grained neural patterns of activation (i.e., patterns with higher spatial frequency). Overall, these results suggest that SND EEG offers potential for future research and clinical use by harnessing significantly more high spatial frequency neural information than current systems.

One strength of our current study is the direct comparison of different ND configurations both against SND and against each other; these comparisons revealed that not only did SND consistently outperform ND, but also uncovered the subtle differences between ND electrode arrangements in neural information capture. Across both the decoding and RSA analyses, there were significant differences in performance for the different ND arrays, indicating that the specific configuration of electrodes in low-density EEG may be important for capturing the most relevant neural activity at the scalp. This, by itself, also indicates that information relevant to classification is present at distances smaller than what spatial Nyquist analysis suggests. Analysis of the “best” and “worst” ND arrays according to the number of electrodes with low signal showed that signal quality was not responsible for the variation in performance across ND arrays (see Supplementary Material). Hence, contrary to spatial Nyquist analysis predictions, ND EEG can lead to suboptimal electrode configurations.

With respect to the different ND electrode configurations, an interesting trend was observed across the different analyses; ND4 had the best ND decoding performance but the worst RSA correlation. This suggests that not only are the specific ND electrode locations on the scalp important, but also that the ideal locations can have different weightings depending on the type of neural information that is being extracted or the analysis being performed (similar to different analyses in fMRI (Nestor, Vettel, & Tarr, 2008)). The fact that SND outperformed ND for all three tests suggests that the additional neural information captured by SND is useful for a wide variety of tasks and analyses, lending support to the idea that SND EEG has significant potential for future research.

As a final point on the potential of SND EEG, we have two reasons to believe that our present results provide a conservative estimate of the extent to which SND improves upon ND in the capture of spatial neural information. First, participants in this experiment had larger-than-average heads (see *Materials and Methods*), and were therefore likely to have thicker skulls, leading to greater decay of high spatial frequency neural information. This implies a participant-bias resulting in lower-than-average signal-to-noise ratio at the scalp for the high spatial frequency neural responses we were attempting to capture. Second, despite very careful gel application, there is also the possibility that neighboring electrodes were sometimes subject to salt bridging, leading to non-independent electrode recordings. Note also, that although it is possible that redundant information from bridged channels might have reduced noise in the SND array, noise reduction cannot be responsible for the pattern of results for two reasons: (1) noise reduction would be expected to improve classification of all six stimuli equally, rather than the observed disproportionate SND advantage for high spatial frequency stimuli, and (2) the EEG RDM measure was normalized for the number of electrodes and therefore revealed similar mean Euclidean distance for the SND and ND arrays, so the higher correlation between SND and the HMAX S1 RDM must be due to more accurate and precise approximations of the neural relationship between stimuli rather than redundant neural information. As such, our results appear to be conservative estimates of the SND advantage and provide a promising foundation for future ultra-high density systems.

Finally, note that although our SND configuration is already a substantial increase in sensor density from current systems, it is likely that further increases in electrode density will result in even greater gains for neural inference, especially with increased trials that enable noise reduction by averaging (Grover & Venkatesh, 2017; Venkatesh & Grover, 2017). In this experiment, the sensor spacing was 14 mm, whereas the standard 10-5 configuration for 128 electrodes has approximately 20-30 mm spacing (Oostenveld & Praamstra, 2001). The inter-electrode distance for current commercial systems is likely to be even greater for people with large head sizes, as was the case in our present experiment. Freeman et al. (2003) suggests that optimal electrode spacing is 10 mm, and recent theoretical estimates indicate that even more neural information can be gathered with sub 5 mm electrode distances (Grover & Venkatesh, 2017). In contrast, the prevailing wisdom, based in part on a spatial Nyquist analysis (Nunez & Srinivasan, 2006) contends that increasing sensor density from current norms (64-128 electrodes) provides minimal benefit. However, here we show that we observe significant increases in the capture of visual neural information by increasing electrode density, particularly for high spatial frequency stimuli. Although the absolute increase in decoding and RSA was modest, ultimately obtaining more neural information at the scalp is also expected to lead to more accurate source localization estimates. Furthermore, engineering SND EEG systems to address particular issues that arise with higher density EEG, such as increased risk of salt bridging, will serve to enhance this SND advantage even more. As such, we conclude that “super-Nyquist” EEG has a great deal of promise for future clinical and neuroscientific research, as well as for brain-computer interfaces. Its full potential should be understood through further experiments with higher density and larger coverage grids that go hand-in-hand with revised fundamental analyses and estimates of limits (Grover & Venkatesh, 2017; Venkatesh & Grover, 2017).

## Acknowledgements

This work was supported by CMU BrainHUB and in part by SRC SONIC center. M. Behrmann was supported by a National Science Foundation grant (BCS-1354350). P.V. was supported in part by the Henry L. Hillman Presidential Fellowship and in part by the Dowd Fellowship from the College of Engineering at Carnegie Mellon University. The authors would like to thank Philip and Marsha Dowd for their financial support and encouragement. M. Boring was supported by a CNBC Undergraduate research fellowship in computational neuroscience. We thank Shawn Kelly and Jeff Weldon for assistance in constructing the EEG cap and Louis (Xiaowei) Kuang for assistance in the decoding analysis.

